# Molecular Characterization of Depression Trait and State

**DOI:** 10.1101/2020.04.24.058610

**Authors:** Rammohan Shukla, Dwight F. Newton, Akiko Sumitomo, Habil Zare, Robert Mccullumsmith, David A. Lewis, Toshifumi Tomoda, Etienne Sibille

## Abstract

Major Depressive disorder (MDD) is a chronic and recurrent brain disorder characterized by episode and remission phases, and poor therapeutic responses. The molecular correlates of MDD have been investigated in case-control settings, but the biological changes associated with trait (regardless of episode/remission) or state (illness phases) remains largely unknown, hence preventing therapeutic opportunities. To address this gap, we generated transcriptome profiles in the subgenual anterior cingulate cortex of MDD subjects who died during a single or recurrent episode or when in remission. We show that biological changes associated with MDD trait (inflammation, immune activation, reduced bioenergetics) are distinct from those associated with MDD phases or state (neuronal structure and function, neurotransmission). On the cell-type level, gene variability in subsets of GABAergic interneurons positive for corticotropin-releasing hormone, somatostatin or vasoactive-intestinal peptide was associated with MDD phases. Applying a probabilistic Bayesian network approach, we next show that gene modules enriched for immune system activation, cytokine response and oxidative stress, may exert causal roles across MDD phases. Finally, using a database of drug-induced transcriptome perturbations, we show that MDD-induced changes in putative causal pathways are antagonized by families of drugs associated with clinical response, including dopaminergic and monoaminergic ligands, and uncover potential novel therapeutic targets. Collectively, these integrative transcriptome analyses provide novel insight into cellular and molecular pathologies associated with trait and state MDD, and a method of drug discovery focused on disease-causing pathways.

**One Sentence Summary:** Integrating transcriptomic with various in-silico analyses identified cellular, molecular and putative biological causal pathways in trait and state depression

## INTRODUCTION

Major depressive disorder (MDD) is the world leading cause of years lost due to disability, with an annual and lifetime prevalence of 6% and 18%, respectively *(1)*. Episodic phases of MDD are characterized by heterogeneous symptoms, including low mood, anhedonia, cognitive impairments, and physiological symptoms (i.e. activity, weight) *(2)*. For the majority of subjects, MDD follows a periodic trajectory of recurring depressive episodes of increasing severity, duration, and of progressive resistance to antidepressants, separated by gradually shortening and incomplete remission phases, leading to treatment-resistance and deteriorating functional fitness (fig.1A) *(3)*. This clinical trajectory suggests the presence of an underlying trait-like and/or progressive neuropathology, which might differ from that of episode or remission “states” of MDD.

**Fig. 1:**
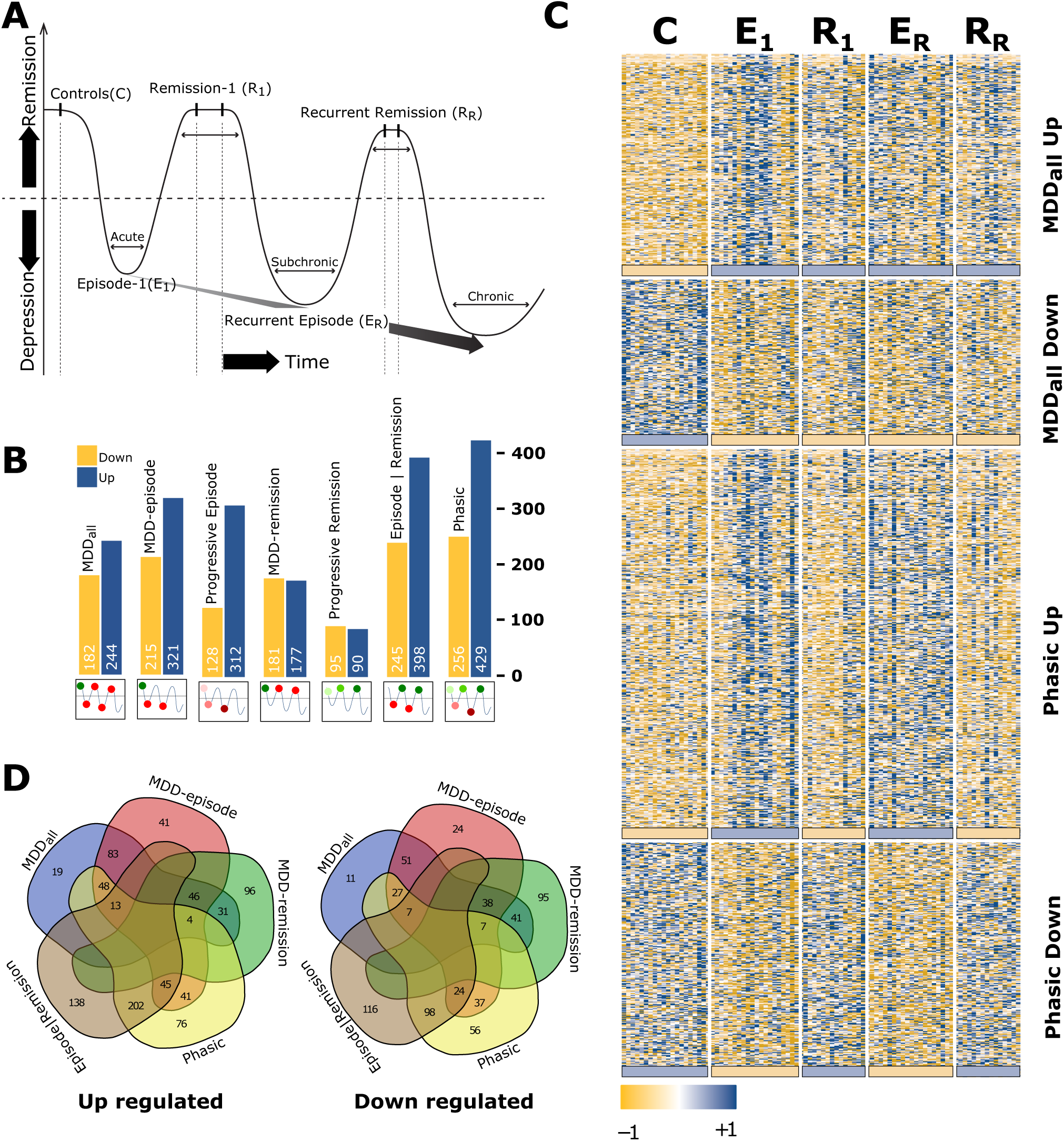
RNAseq-based identification of state-dependent and phasic molecular changes in MDD. **A)** Clinical evidence frequently shows recurrent episodes of MDD (valleys) of increasing severity, reduced therapeutic response and shorter remission periods (crests), suggesting the presence of a progressive underlying pathology across phases (black arrow). **B)** Numbers of differentially expressed genes associated with different MDD phase contrasts (p < 0.05). The text (top) and graphics (bottom) describe the contrasts used in the text and figures. **MDD-all**: all MDD cohorts versus Controls. **MDD-episode:** two MDD episode cohorts versus Controls (E_n_|C). **MDD-remission:** two remission cohorts versus controls. **Episode/Remission:** two MDD episode cohorts versus two remission cohorts (E_n_|R_n_). **MDD-phasic**: capturing the phasic up- and downregulated gene expression changes correlated with episode and remission phases. **Progressive Episode**: monotonic increase or decrease across control and episode groups; C→E_1_→R_R_. **Progressive Remission**: monotonic increase or decrease across controls and remission groups; C→R_1_→R_R_). **C)** Heatmap of gene expression changes corresponding to the MDD**-all** (top 2 panels) and **MDD-phasic** (bottom two panels) contrasts. The colored bars at the bottom of each panel are visual summaries of gene effects within the section above. The MDD-all contrast shows consistent up or downregulation of genes across the various phases of MDD, compared to controls. The MDD-phasic contrast shows “waves” of gene changes that coincide with the episode and remission phases of MDD. **D)** Venn Diagram showing intersection of up- and down-regulated genes associated with MDD-all, MDD-episode, MDD-remission, Episode|remission and Phasic contrast. Note the highest intersection between Episode|Remission and Phasic contrasts.

The subgenual anterior cingulate cortex (sgACC) lies at the intersection of bottom-up sensory input and top-down cortical control and is involved in integrated processing of emotions, including mood and reward. The activity of sgACC is increased in subjects with MDD, as well as in individuals with high neuroticism, fear of peer rejection and healthy humans during experimentally induced sadness *(4–6)*. An effective antidepressant therapy reverses the hyperactivity of the sgACC, making it the target and suggested mediator for therapeutic effect of deep brain stimulation *(7, 8)*. Magnetic resonance spectroscopy and transcranial magnetic stimulation studies suggest reduced gamma-aminobutyric acid (GABA) levels and cortical inhibition as a mechanism of sgACC dysregulation in MDD, which too appear to normalize after successful treatment *(9–12)*. Large-scale transcriptomic studies in MDD postmortem sgACC samples demonstrate dysregulation in cytoskeleton, rearrangement of neuronal processes, synaptic function and presynaptic neurotransmission *(13, 14)* associated with GABA and glutamate receptor signaling. At the cellular level, reduced glial and increased neuronal densities were reported, associated with reduction in axon and dendrites *(15, 16)*. Reduced expression of markers for GABAergic interneurons targeting either the dendritic (e.g. somatostatin) or perisomatic (parvalbumin) compartments of pyramidal cells were reported, associated with reduced neurotrophic support *(17–19)*.

To go beyond case-control studies and investigate molecular correlates of disease trait or state, we recently performed a large-scale proteomic study in a postmortem cohort of subjects in first or recurrent phase of episodes or remission, and in healthy controls *(20)*. In addition to state-specific results, this study highlighted robust MDD trait-associated changes (i.e., regardless of episode or remission state), suggesting a continuous underlying pathology *(20)*. However, limitations in detection and sensitivity of the proteomic approach precluded a full molecular characterization of the illness phases and putative progression. In contrast, RNAseq-based transcriptome analysis provides a broader snapshot of functional state and transcriptome-based systems biology analysis have proven appropriate for investigating complex diseases and putative causal pathways *(21–23)*.

Here we hypothesize that RNAseq-based transcriptome profiling of human postmortem sgACC samples from patients in depressive or remission phases of MDD and in controls, combined with ontological, systems biology, Bayesian network and perturbagen-induced transcriptome analyses, would enable the molecular characterization of disease trait, state and chronicity, as well as putative causal biological changes and therapeutic targets.

## RESULTS

### MDD trait and state are characterized by distinct biological changes in sgACC

RNAseq-based expression profiles from sgACC samples were obtained from one control (C) and four cohorts of MDD subjects during a first episode (E_1_), remission after first episode (R_1_), recurrent episode (E_R_) or remission after recurrent episodes (R_R_) (fig.1A, table S1). Differential expression and biological pathway analyses were performed along the following group contrasts (fig.1A and Methods): 1)**MDD-all**: all MDD cohorts versus Controls, 2) **MDD-episode:** two MDD episode cohorts versus Controls, 3) **MDD-remission:** two remission cohorts versus controls, 4) **Episode/Remission:** two MDD episode cohorts versus two remission cohorts, 5) **MDD Progressive Episode**: monotonic increase or decrease across control and episode groups, 6) **MDD Progressive Remission**: monotonic increase or decrease across controls and remission groups; 7) **MDD-phasic**: capturing gene expression patterns coupled to the phasic changes across groups.

The differential expression analysis revealed distinct gene sets with expression changes matching the illness trait and state with the following features: Overall, upregulated genes were more common than downregulated genes across contrasts (fig.1B, table S2); The **MDD-all** contrast, representing trait, showed consistent up- or down-regulation patterns of gene expression regardless of episodes or remission phases compared to controls (fig.1C, top two panels); The **MDD-phasic** contrast revealed robust patterns of gene expression that oscillated with episodes and remission phases (see “waves” in bottom two panels in fig.1C). The MDD-phasic gene set overlapped substantially with the direct **Episode/Remission** contrasts (fig.1D), which captured the range of the “wave” pattern; Minimal overlap was observed between gene sets identified in the **MDD-episode** and **MDD-remission** contrasts (fig.1D) representing MDD states, or between the **Progressive-episode** and **Progressive-remission** contrasts (less than 1%).

To summarize the vast data, gene expression changes across different contrasts were next analyzed at the level of biological pathways (p-value <0.01; FDR<0.25) and functional themes. Consistent with gene expression findings, upregulated pathways were more common than down-regulated pathways (fig.2, Table S3). For the **<u>MDD-all</u>** <u>contrast</u> defining the illness trait, multiple pathways associated with inflammation, immune system, angiogenesis and vascular growth factors were found to be upregulated. Other upregulated pathways included transcription associated with stress response and multiple aspects of protein modification processes (e.g., involving *JNK cascade, MAPK activity, auto-ubiquitination*; table S3). Few down-regulated pathways were observed and were associated with cellular bioenergetics (*Mitochondrial translation associated process, mitochondrial activity, ATP metabolic process)* and with *neuropeptide signalling*, including signalling and cell-type gene markers.

**Fig. 2:**
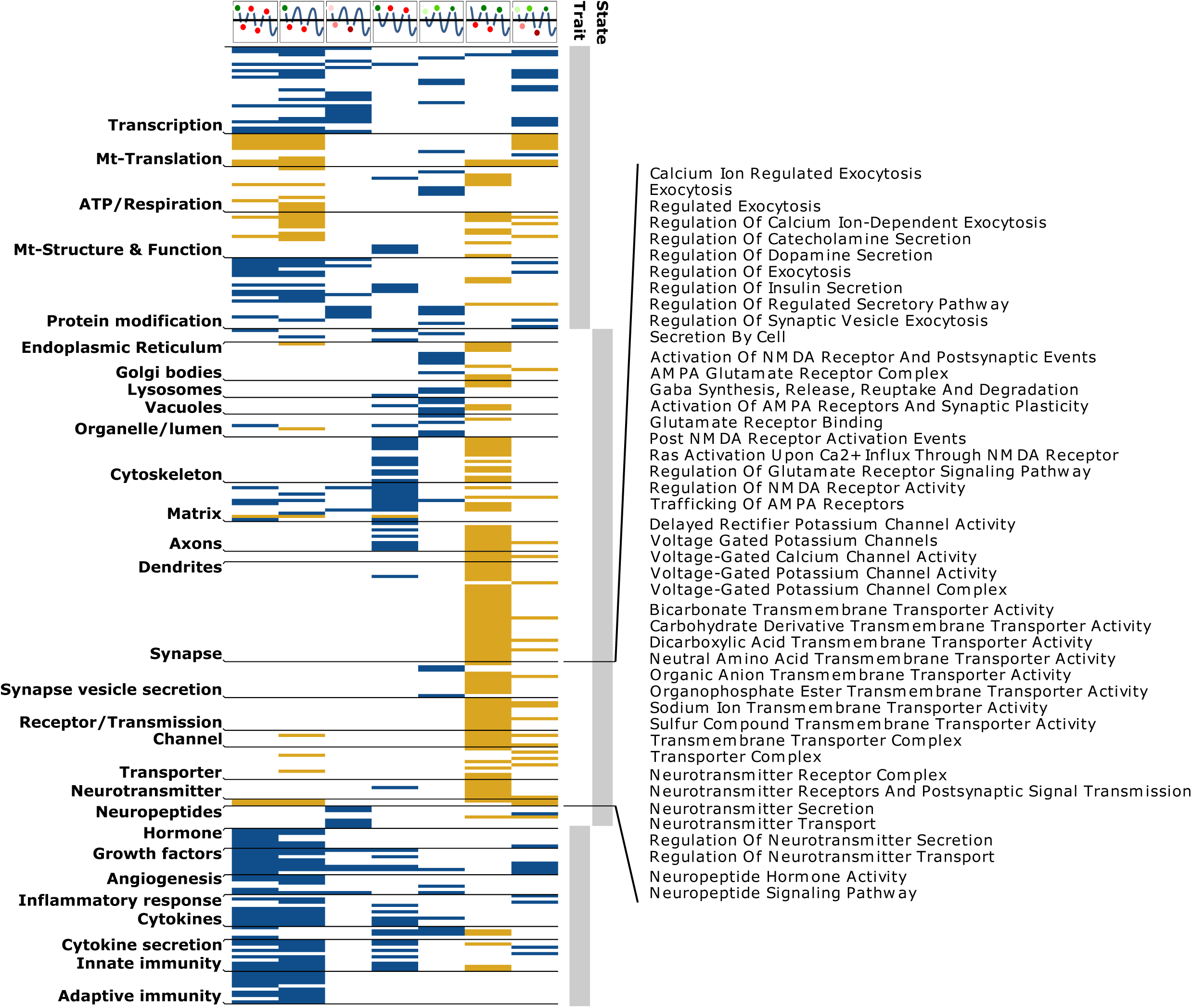
Profiles of biological pathway affected in the MDD group contrast and validation studies. Altered pathways associated with different biological themes (left) per contrast. The blue and yellow colors represent up and down regulated pathways scaled by enrichment scores, ranging from +1 (Blue) to -1 (Orange).

Results for the **<u>MDD-episode</u>** <u>contrast</u> were highly similar to **MDD-all** contrast (fig.2, Columns 1-2), including upregulated immune and inflammation-related pathways, upregulated translation related to stress response and aspect of protein modifications, as well as downregulated pathways related to mitochondrial bioenergetic functions. For neurotransmission, pathways associated with *Voltage gated potassium channels* and *transporter* activities were downregulated, in addition to the same prior neuropeptide signaling pathways. The **Progressive-episode** contrast was more selectively associated with upregulated pathways related to transcription, protein modifications, angiogenesis and immune system, suggesting increasing pathological changes related to these pathways with successive MDD episodes. Interestingly, several pathways related to cellular responses to hormones were selectively upregulated in this contrast.

The **<u>MDD</u>**<u>-</u>**<u>Remission</u>** <u>contrast,</u> similar to the MDD-all and MDD-episode contrasts, was associated with upregulated inflammation- and immune-related pathways, although immune-related pathways were limited to innate versus both adaptive and innate immunity in the Episode contrast (fig.2, Columns 1-3). Notably, changes more specific to the **MDD-Remission** contrast included multiple upregulated pathways related to cytoskeleton, axonal and extracellular matrix, suggesting a structural cellular reorganization during remission phases of the illness. Changes associated with the **Progressive remission** contrast were associated with upregulated *bioenergetic and cellular respiration* pathways, upregulated cellular *organelle activity (Endoplasmic reticulum, Golgi bodies, Lysosomes Vacuoles and lumen, secretion)* (Fig.2), and with few pathways associated with inflammation and the immune system.

Finally, the direct **<u>Episode/Remission contrast</u>** was associated with downregulated bioenergetic-related pathways (*ATP, cellular respiration*, and *Mitochondrion-related* pathways), consistent with findings form the Episode- or Remission-contrasts. Surprisingly, we observed an additional massive downregulation of multiple pathways related to the structure and function of all neuronal compartments (*cytoskeleton, organelles, matrix, axon, dendrites synapse, channel, receptor/transmission dendrite, vesicle secretion, extracellular matrix)*. These changes included pathways associated with multiple aspects of synaptic function and neurotransmission, such as downregulated neurotransmission mediated by glutamate, GABA, dopamine, catecholamines potassium and calcium voltage-gated channels, and multiple membrane transporter functions. This suggests the presence of multiple moderate state-dependent changes along different phases of episode and remission, affecting multiple aspects neuronal signaling, which are only detectable by direct comparison across opposite phases, and that are missed here when comparing MDD subjects to control groups. Indeed, an analysis for changes matching the oscillatory nature of the disease trajectory (**MDD-phasic**) identified most functional themes of the Episode/Remission contrast (fig.2, Column 6), although to lower degrees than separate contrasts.

Together the results of the various group contrasts demonstrate the presence of two parallel pathological entities, one corresponding to a depressive trait, generally associated with inflammation, immune system activation and decreased bioenergetics, and a second state-dependent pathological entity matching the clinical phases of the illness, and associated with changes in the structure and function of cells, and with multiple aspects of neurotransmission, both generally negatively impacted during depressive states.

### Validation of altered gene expression and pathway profiles

Results were validated using three approaches. First, a comparison to results obtained in a similar study performed by Pantazatos et al *(24)*, using RNAseq in dorsolateral prefrontal cortex of MDD and control samples, revealed significant similarities in gene expression changes in that study with the MDD-episode (p-value=0.05), MDD-phasic (p-value=0.046) and Progressive episode (p-value=0.043) contrasts, and trend-level overlap of MDD-all (p-value=0.062). Second, we compared the current RNAseq results with previous mass spectrometry-based proteomics results in the same cohort *(20)*. Gene and protein expression significantly correlated (n=3000 genes/proteins; R=0.34; p=9.34×10^−94^), and significant overlaps between protein and gene changes were observed for the MDD-episode (p-value=0.02), Episode/Remission (p-value=0.043), and trend-level overlap for the MDD-remission (p-value=0.062) contrasts. Third, technical validation was performed by qPCR for selected genes among affected pathways within different contrasts (fig.3A). qPCR and RNAseq results were highly concordant (fig.3B).

**Fig. 3:**
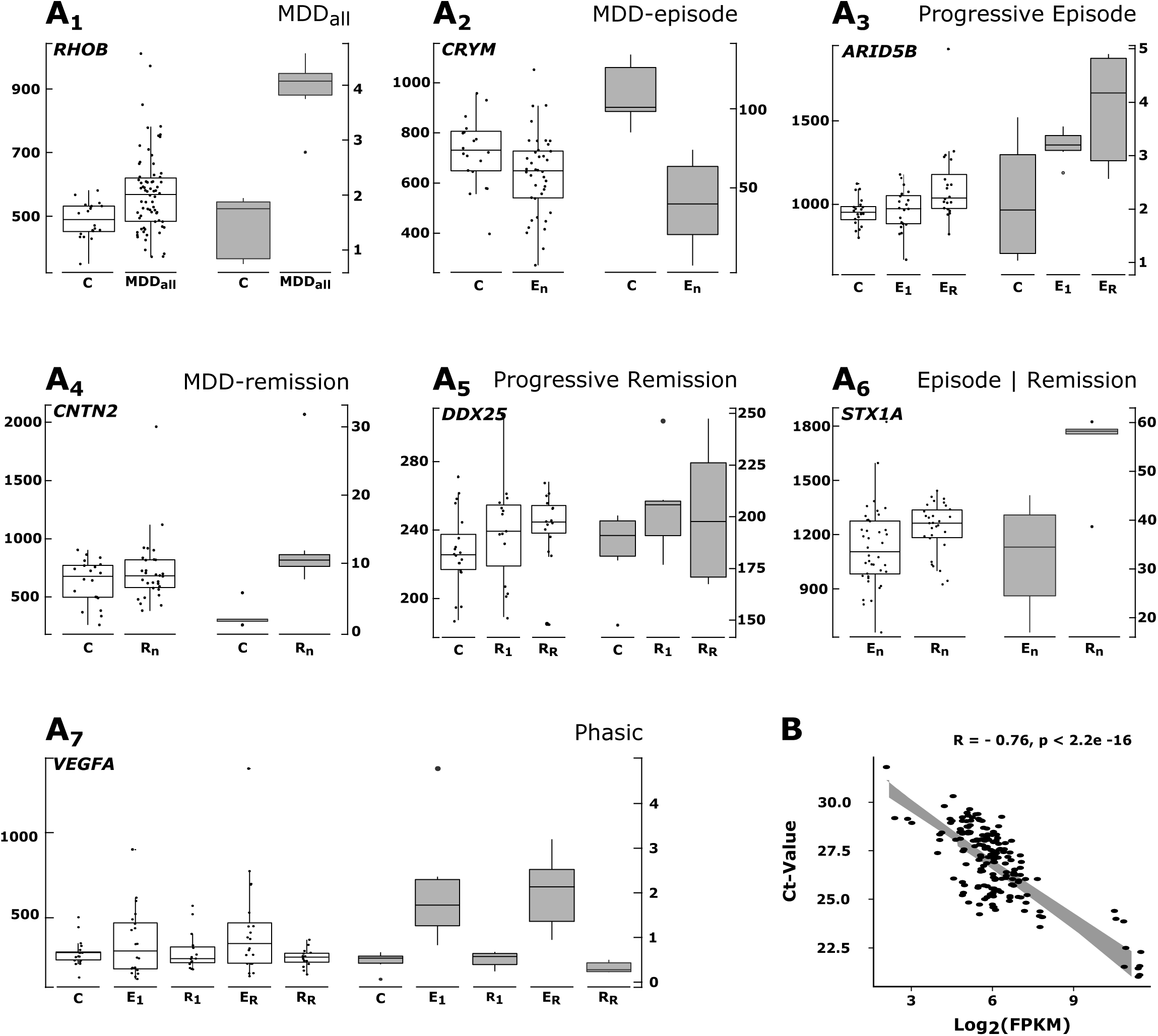
qPCR validation of differentially expressed genes and representative of affected pathways in each contrast. In each image (from **A1-A7**) left white box plots represent RNAseq data. Right grey box plots represent qPCR data (n=5/group) **A1) *RHOB***: belonging to GO group ‘angiogenesis’ and up regulated in all MDD subjects compared to control in MDD-all contrast. **A2) *CRYM:*** belonging to GO group ‘mitochondria’ and ‘oxidation-reduction process’ and down-regulated in Episodic subjects compared to control in MDD-episode contrast. **A3) *ARID5B:*** belonging to GO group ‘regulation of transcription, DNA-templated’ and progressively Up-regulated in episodic states. **A4) *CNTN2:*** belonging to GO group ‘microtubule cytoskeleton organization’ and Up-regulated in Remission subjects compared to control in MDD-remission contrast. **A5) *DDX25:*** belonging to GO group ‘regulation of translation’ and progressively Upregulated in remission states. **A6) *STX1A:*** belonging to GO group ‘presynaptic membrane’ and Upregulated in Remission subjects compared to episodic subject in Episode|Remission contrast. **A7) *VEGFA:*** belonging to GO group ‘angiogenesis’ and coupled with phasic changes involved in depression. P-values < 0.05 for all independent RNAseq and qPCR group comparisons. B) Correlation between RNAseq and qPCR was calculated for all the genes selected for validation (**A1-A7**). Data from all the gene were pooled and each point in the correlation plot represents one subject.

### RNA-seq data deconvolution identified a subpopulation of interneurons affected by phasic state changes in MDD

Our present data was derived from gray matter samples where relative cell proportions and associated biological signals are masked. To address this limitation, we used existing layer-specific single nucleus RNAseq data from ACC (from Allen brain atlas) and statistical methods to deconvolute the data. We identified 19 cell-type clusters (fig.4A), which we characterized in two stages: First, globally, based on known markers associated with pyramidal neurons (SLC17A4), interneurons (GAD1 or GAD2), astrocytes (GFAP), oligodendrocytes (OPALIN), microglia (CX3CR1) and neuroglia (PDGFRA) (fig.3B), and second, locally, based on the genes most enriched in each clusters (fig.4C). For instance, the GABAergic interneuron markers PVALB, SST, VIP and CRH were amongst the most enriched genes in cluster 5, 4, 7 and 8 respectively. Next, we identified sets of genes that clearly discriminated each of the 19 cell-type clusters (fig.4D, table S4). These discriminative gene sets were then used to estimate the relative proportion of each cell type in the MDD datasets obtained from each cohort, using a support vector machine (SVM) approach *(25)*. Results show that cluster 8 displayed lower expression of its gene markers in episode compared to remission phases (p<0.042, fig.4E, right panel), and that these changes followed a phasic trajectory between single and recurrent phases of the illness (p<0.016, fig.4E, left panel). Cluster 8 corresponds to GABAergic interneurons expressing the SST, VIP and CRH gene markers (fig.4C). Notably, these markers correspond to the neuropeptide signaling GO group previously identified in the biological pathway analysis of the MDD-All, MDD-episode and MDD-Phasic contrasts (Fig.2). An analysis with layer-specific single cell data further showed an enrichment of cluster 8 gene sets in in cortical layers 1 and 2/3 (fig.3F).

**Fig. 4:**
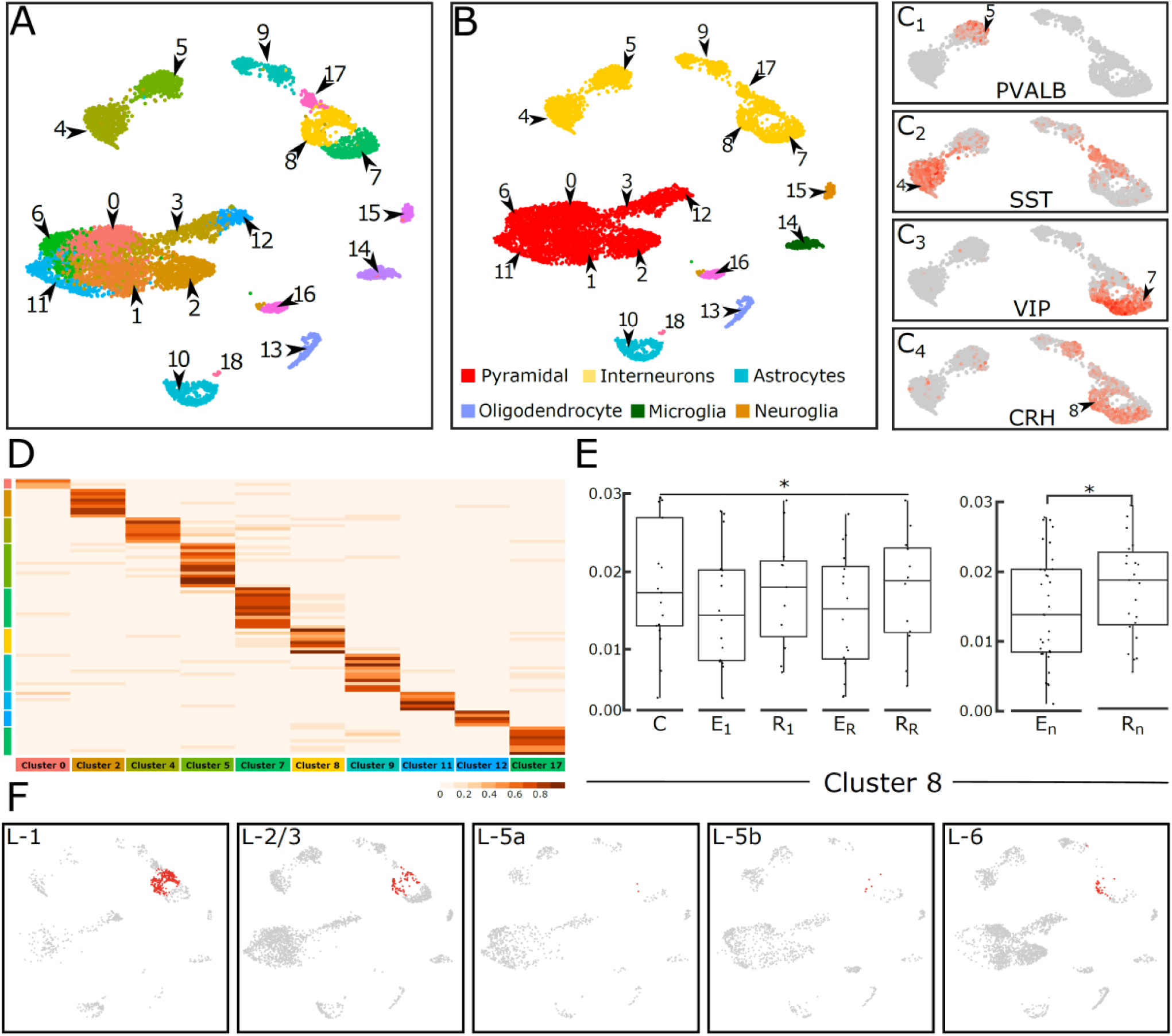
Cell-type deconvolution of gray matter RNAseq reveals gene expression changes in synchrony with MDD phases for CRH-, SST- and VIP-positive GABAergic interneurons. **A)** Clustering of human anterior cingulate cortex single nuclei RNAseq data from Allen Brain Atlas. **B)** Clusters were characterized globally using known markers of Pyramidal neurons, interneurons, astrocytes, Oligodendrocytes, microglia and polydendrocytes (see results section for markers). **C)** Clusters were characterizing locally using the top enriched expressed genes (upregulated) by cluster. **C**_**1**_) Cluster 5 enriched with PVALB. **C**_**2**_) Cluster 4 enriched with SST marker. **C**_**3**_) Cluster 7 enriched with VIP. **C**_**4**_) Cluster 8 enriched with CRH. Note that some clusters have overlapping expression of different markers. For instance, cluster 8 (CRH) is also enriched in SST and VIP. **D)** Heatmap of highly discriminative gene, i.e. markers used as reference for estimating relative cell proportions in **(E)**. Rows represent genes and columns represent the cluster identified in (A). Only representative clusters are shown here. For a heatmap of all the clusters see supplementary fig.X. **E)** Proportion differences of cluster 8 cell types in Episode/remission contrast (top) and phasic contrast (bottom). The y-axis shows the relative proportion of cluster 8 cell types while the x-axis represents different cohort used in the study **F)** Layer specific distribution of cluster 8 neurons (yellow). Note the high enrichment of cluster 8 neurons in supragranular layer 1 and 2/3.

We next sought to validate these findings using gene expression datasets obtained in independent cohorts. We used a list of 566 genes obtained in a meta-analysis of altered gene expression in MDD across corticolimbic regions, including the sgACC *(26)* (table S5) and performed a hypergeometric overlap analysis in all 19 gene clusters (table S4). Within the interneuron clusters (Yellow in Fig 4B), the meta-analysis gene set overlapped significantly with the previously identified cluster 8 (q-value < 0.03), corresponding to interneurons expressing CRH, VIP and SST, hence validating the current results.

Next, we investigated the putative impact of altered Cluster 8 cell types (fig.4A-C) on overall biological changes. For this we focused on the most significant contrast (**Episode/Remission)** and compared gene expression differences with or without regressing out the variability associated with the putative cell-type proportions differences of cluster 8 (See Discussion). 721 (Up: 419, Down: 302 with respect to Remission) genes showed lower p-values after regression (table. S6). These combined gene changes corresponded to the following downregulated biological pathways in episode: *voltage-gated potassium channel complex* (q<1.06×10^−2^), *postsynaptic membrane* (q<2.7×10^−3^), *asymmetric synapse* (q<3.88×10^−3^), *axon-ensheathment* (q<4.8×10^−3^) and *regulation of cell morphogenesis* (q<2.21×10^−2^), and no upregulated pathways (table. S6), consistent with the analysis of biological pathways associated with the Episode/Remission contrast (Fig. 2A, insert). Finally, to assess cluster 8 specific differential expression, we included the interaction between cluster 8 cell proportion and variables (i.e. Episode and remission state) of **Episode/Remission** contrast in a statistical linear model *(27)* we identified 75 upregulated and 175 downregulated (table. S7, p-value<0.01), corresponding to upregulated *chemokine binding* (q-value < 0.04) and downregulated *nf-KappaB* (q-value < 0.01) pathways.

Taken together, inferring cell-type origin of gene expression revealed state-dependent phasic biological changes affecting upper layers GABAergic interneurons identified by the SST, VIP and CRH cell-type markers. Moreover, the data suggest that these cell type-dependent changes occur in synchrony with the broader reduced neuronal signaling phenotype observed in the MDD Episode/Remission contrast and may be directly affected by inflammation/immune-related events.

### Investigating causality using Bayesian gene network analysis

Whereas the previous group contrasts and cell-based analyses provide biological and cellular insight into MDD state and phases, they do not inform on causal links. To identify putative causal pathways, we summarized the expression profiles across all five cohorts into modules of co-expressed genes using consensus weighted gene co-expression network analysis. This analysis reduces expression profiles across all five cohorts into common, more stable, and coherent functional modules of co-expressed genes that are comparable across all cohorts. The first principal component summarizing omnibus gene expression in each module (i.e. eigengene) was used to construct a Bayesian network. A Bayesian network models the probabilistic dependencies between gene modules and with the disease node, and is represented as a directed acyclic graph (DAG)*(28–30)*, where edges represent the probabilistic dependencies between gene modules, organized from early to later module (referred as parent to child).

22 consensus modules were identified, containing from 31 to 4494 genes. 18 of these modules were significantly enriched for at least one gene ontology (GO) functional category at q-value < 0.05 (fig.5A, table. S8) and were used to create the DAG (fig.5B). The direction of the edges and probabilistic dependencies between modules were first inspected for biological and theoretical consistency. For instance, module M05, the initial source of all modules, is enriched in pathways associated with *system development* (q-value<1.52×10^−9^) and connected to module M02 and M011, enriched in pathways associated with *nervous system development* (q-value<4.03×10^−11^) and *anatomical structure morphogenesis* (q-value<6.5×10^−3^), respectively. Similarly, the known relationship between the immune system and cytokine *(31)* is captured by the high probability parent-child association between module M21, enriched with immune response (q-value<8.67×10^−3^), and module M10, enriched with pathways associated with cytokine response (q-value<4.79×10^−27^).

**Fig. 5:**
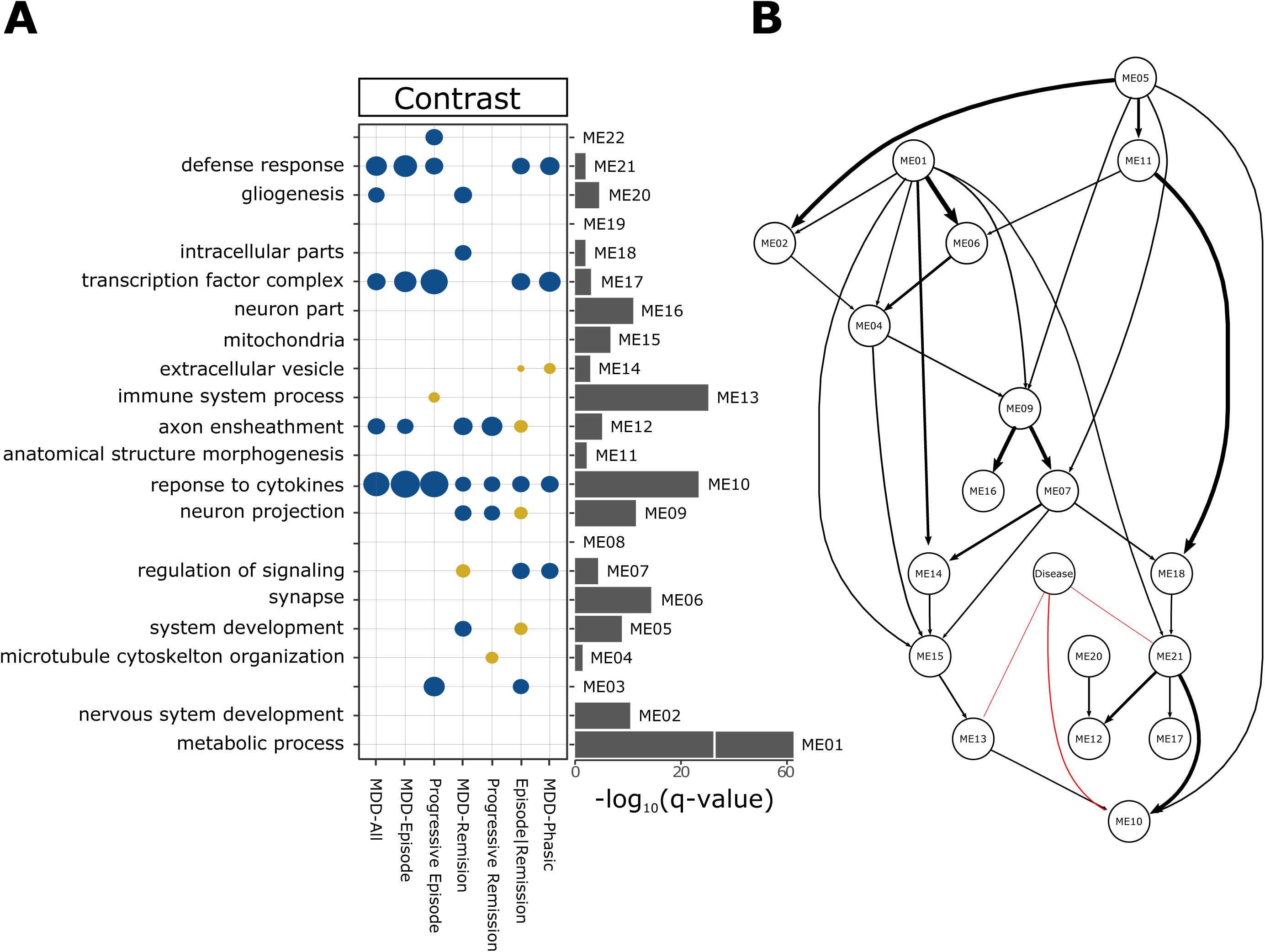
Prioritizing putative causal gene modules in MDD using Bayesian network: **A)** Consensus gene modules were identified using the combined RNAseq data across diseases phases and characterized using GO enrichment. GO terms highly enriched for each module are shown on the left, and -log10 p-values (fdr <0.05) of enrichment tests for the GO term are shown on the right. The middle panels show the enrichment of modules in the MDD contrasts (From Fig.1B). The blue and yellow circle shown enrichment in up- and down-regulated genes for each module. The size of circles is proportional to the -log10 of p-value associated the enrichment. Note that module ME03, ME08, ME19 and ME22 were not associated with any GO terms and were removed from further analysis. **B)** Directed acyclic graph (DAG) obtained after fitting Bayesian network to the eigengenes of characterized consensus modules, shown as nodes. The thickness of the edges connecting the nodes is proportional to the number of times, in percent, the edge was detected in 1000 permutation used to fit the Bayesian network. The connection of “Disease” node (methods) to its child nodes are shown in red.

The MDD node is linked to three DAG modules: module M13, enriched with *innate immune response* (q-value<1.60×10^−17^), module M10, enriched with *cytokine response* (q-value< 1.31×10^−21^), and *response to stress* (q-value<1.72×10^−17^), and module M21, associated with *defense response* (q-value<1.04×10^−2^) and *stress response* (q-value<4.63×10^−3^) mostly associated with oxidative stress, as suggested by the child module M17, enriched with *response to oxidative stress* (q-value<4.6×10^−3^) and *cellular response to stress* (q-value<2.05×10^−3^, table. S8). Among the 3 MDD-associated modules, M10 had the highest probabilistic association to the disease module and is a child to disease-associated M13 and M21, making it the endpoint of the graph and the most disease-associated module. This is supported by the enrichment of M10 with upregulated pathways in all seven contrasts outlined above (fig.5A).

We next investigated the hypergeometric overlap of discriminant markers of the 19 cell-types clusters (fig.4D) with the MDD-associated modules. Module M10 was enriched in astrocyte (cluster 10, p < 0.02) and microglia (cluster 14, p < 5.3 × 10^−9^); module M13, in SST neurons (cluster 04, p < 0.03) and microglia (cluster 14, p < 5.55 × 10^−127^) and module M21 was enriched in none of the cell markers. As an independent validation, we looked for hypergeometric overlap with cell specific gene set available from a previous study *(32)*. Module M10 was enriched in microglia (p < 0.01), module M13 in microglia (p < 2.23 × 10^−27^) and module M21 in endothelial (p < 0.03). Overlap of module with lamina-specific gene sets *(33)* suggest the enrichment of all three modules in layer 1 (M10: p < 1.2 × 10^−14^; M13: p < 2.29 × 10^−27^; M21: p < 0.03) which also coincides with the significant enrichment of disease associated cluster 8 cells in superficial layer 1 (Figure 4F).

Together, the Bayesian network analysis organized the biological variability observed across the control and MDD cohorts (as measured by gene coexpression) into a coherent and directional graph. While these biological events occur in the context of the various clinical phases of MDD, the direct association of the disease module with three DAG modules suggests that immune and cellular stress functions associated with these modules are more proximal, and potentially causal, to the trajectory and clinical manifestation of the illness. Moreover, cell-enrichment analyses implicate superficial layer inhibitory neurons, glial and immune cells as potential mediators of putative causal biological events in MDD.

### The MDD gene module identified in the Bayesian network is associated with known antidepressant drugs related pathways and novel drug targets

Based on the assumption that the MDD-associated DAG end-module M10 may have a causal role in the illness, we next investigated whether the M10 gene expression profile could be mimicked or antagonized by drugs. For this we probed the M10 associated gene-set against the connectivity map (cmap) database, a catalog of transcriptomic responses to multiple molecules in diverse cell types. Despite many limitations of the interventions (simplified cell systems, single doses, acute drug exposure, few neuronal cell lines), the use of the cmap database has been instrumental in identifying novel therapeutic modalities in complex diseases *(34)*, including in neurology and neuropsychiatry *(35)*.

33 and 50 molecules were identified as antagonizing (fig.6) or mimicking (fig.7) the M10 gene expression profiles in neuronal cell lines, which, based on the signature reversion principal *(34)*, may reflect therapeutic or MDD-inducing effects, respectively. Among the 33 M10-antagonizing molecules, three were either dopamine, its precursor (glutamyl-dopamine) or its structural variant (6-nitrodopamine), 15 (45.45%) have G coupled protein receptors (GPCRs) as their target class, of those 5 were dopaminergic, 4 were serotonergic receptors, and two targeted the serotonin and/or dopamine transporters. The other major class of the drug target includes enzyme (6/33), electrochemical transporter (4/33) and nuclear receptors (3/33). Notably, 7 of the identified drugs are either clinically used to treat neuropsychiatric disorders or identified as a potential drug in human postmortem studies (spermidine) *(36)*.

**Fig. 6:**
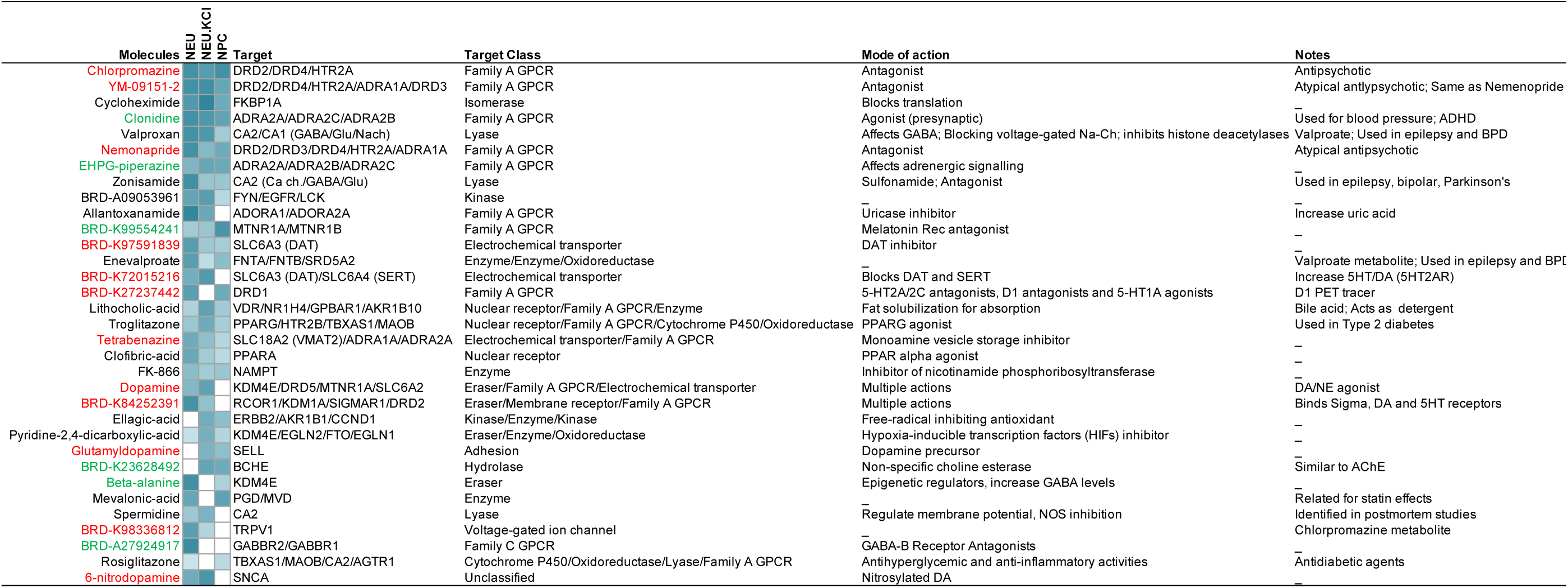
Connectivity mapping of antagonizing molecules: Dissimilarity (antagonizing effect), represented by tau (see methods), of drug-induced transcriptomic profiles and module 10 gene sets is shown. Drug-induced profiles are derived neuronal cell lines. Significant Tau values of <-90 are shown in increasing shades of blue. Molecules in red are associated with either monoamine or catecholamine transmitter while those in green with other known neurotransmitter. The target class column represents predicted protein targets using SwissTargetPrediction. Mode of action and notes taken from pubchem

Among the 50 MDD-mimicking molecules (fig. 7), 12 (24%) were of GPCRs target class, of those 6 were serotonin and 3 were dopamine receptors (mostly with opposing effects than compounds antagonizing the M10 expression profile), and one each for opioid, melatonin and adrenergic receptors. The other major class (> 10% of total) includes eraser (9/50, i.e. HDAC inhibitors) and protease (5/50).

**Fig. 7:**
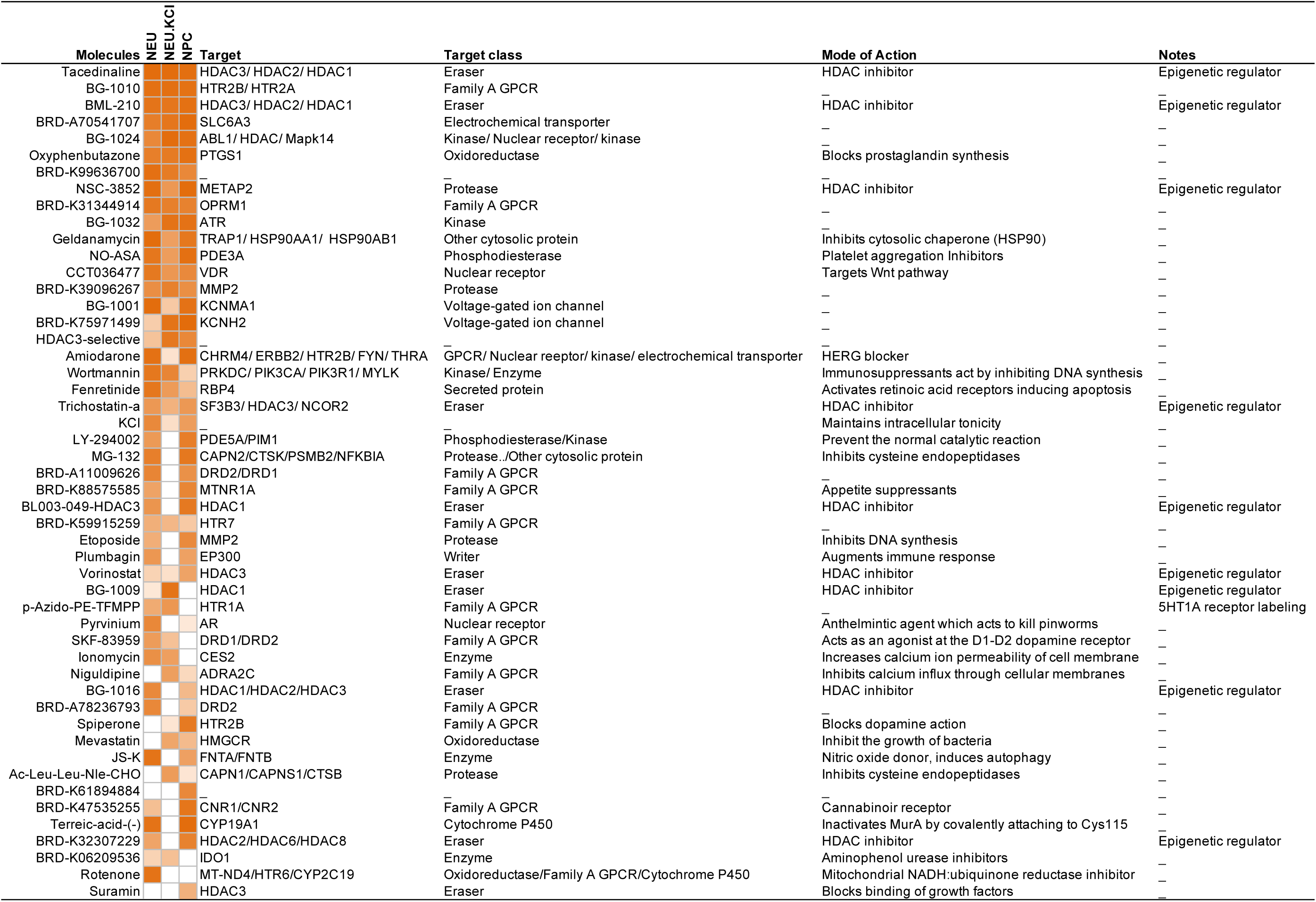
Connectivity mapping of disease mimicking molecules: Similarity (**disease mimicking effect)**, represented by tau (see methods), of drug-induced transcriptomic profiles and module 10 gene sets is shown. Drug-induced profiles are derived neuronal cell lines. Significant Tau values of >90 are shown in increasing shades of orange. The target class column represents predicted protein targets using SwissTargetPrediction. Mode of action and notes taken from pubchem.

In contrast, using the M16 gene module, which is part of the DAG, but not enriched in any of the MDD contrasts (fig.5A), only 7 and 9 molecules were identified with antagonizing and mimicking effects, respectively and the target class associated antagonizing molecules belonged to Eraser (3/7), nuclear receptor (2/7), kinase (1/7) and voltage gated channel (1/7).

## DISCUSSION

Unraveling the complex biological disturbances occurring in heterogeneous brain disorders, such as MDD, is a major challenge that is further complicated by the frequent trajectory of recurrent episode and remission phases of the illness. This symptomatic profile suggests that pathophysiological disturbances underlying MDD may wax and wane, however this contradicts the other clinical observation of progressive development of treatment-resistance and deteriorating functional fitness, suggesting a continuous and progressive pathophysiology. Here, using RNAseq-based transcriptome profiles from human postmortem sgACC samples obtained from MDD patients who died during depressive or remission phases, and from control subjects, we show the presence of two parallel pathological entities (fig. 1B). The first pathology corresponds to a stable MDD state, regardless of episode and remission and is associated with inflammation, immune activation, and reduced bioenergetics. The second is a state-dependent pathology, which consists of dynamic biological changes that match the clinical phases of the illness and that are mostly detected when directly comparing the transcriptomes of episode versus remission subjects. This state-related pathology is associated with neuronal structure and function, affecting both fast neurotransmitter (GABA, Glutamate) and neuromodulator (catecholamines, neuropeptides) systems, sets of changes that are downregulated during episodes and not observed or reversed during remission phases. Although these studies were performed using combined cortical gray matter and in cross-sectional cohorts, two additional *in silico* analyses provide perspective on cellular specificity and putative causal processes. First, cellular deconvolution approaches showed that variability in gene sets associated with GABAergic interneurons positive for corticotropin-releasing hormone, somatostatin or vasoactive-intestinal peptide was associated with MDD phases. Second, a probabilistic Bayesian network approach showed that gene modules enriched for immune system activation, cytokine response and oxidative stress, may exert causal roles across MDD phases. Finally, using a database of drug-induced transcriptome perturbations, we show that MDD-induced changes in putative causal pathways are antagonized by families of drugs associated with clinical response, including dopaminergic and monoaminergic ligands, and suggest an innovative approach to uncover potential new therapeutic targets. Collectively, these integrative transcriptome analyses provide novel insight into cellular and molecular pathologies associated with trait and state MDD, and a method of drug discovery focused on disease-causing pathways.

### Dissociating trait and state pathologies of MDD

The identification of two parallel pathological entities corresponding to trait and state MDD is graphically represented in the Figure 2 cluster graph, and supported by coherent results from the analysis of biological pathways associated with these patterns. Starting with the MDD trait pathology, the results show a robust link to inflammation and immune activation, consistent with prior reports *(41)*. These events often recruit extensive signal transduction pathway protein modifications, such as phosphorylation of proteins in the MAPK pathway (ref), and increased demand on local blood supply, mediated by increased angiogenesis, which are both also associated here with MDD trait pathology. These findings are consistent with a large body of literature on MDD for inflammation *(39)*, immune system activation *(40)*, MAPK activation (ref), but less so for angiogenesis, as increasing angiogenesis has been proposed as a therapeutic avenue (ref).

Note that “trait” is defined based on statistical association with the MDD-All contrast, i.e. subjects in episode or remission phases of the illness compared to controls, but some of the identified trait phenotypes displayed various degrees of significance in the separate contrasts. For instance, pathways related to innate immunity were consistently upregulated in all three contrasts, but changes in adaptive immunity and inflammatory response were limited to the MDD-All and MDD-episode, but not in the remission state, suggesting additional phasic state-dependent immune features. Reduced bioenergetics was identified as trait pathology, represented by decreased expression of genes implicated in mitochondrial structure and function, and in cellular energy production (i.e. ATP). This was observed in the MDD-All and MDD-Episode contrasts, but not the MDD-remission, suggesting a partial reversal of the phenotype during remission, as supported by significant results in the direct Episode/remission and phasic contrast analyses.

The MDD state pathology (as identified in group contrasts, but not the MDD-all contrast) was associated with classical neurotransmission changes often reported in depression (REFS), negatively affecting excitatory glutamatergic (NMDA and AMPA receptors), inhibitory GABAergic systems, multiple aspects of voltage-gated channel activities, and with reduced structural elements of neurotransmission (axons, dendrites, vesicle machinery, etc), consistent with anatomical studies showing reduced pyramidal neurons dendrites and sizes (Refs). The fact that these results are only observed when comparing opposite groups (i.e. Episode/Remission contrast) suggests moderate state-dependent changes affecting neuronal signaling that are more difficult to identify in traditional case-control studies and that would be more sensitive to cohort composition and sizes, potentially explaining past discrepancies across studies (refs). In contrast, neuronal structural components (cytoskeleton, matrix, axons) were restored during remission phases (i.e. significant in the MDD-Remission contrast), and a progressive upregulation of intracellular vesicles associated with trafficking, and lysosomal and autophagy activities, was observed with successive episodes (i.e. Progressive-Remission contrast). These results show that the often-reported neurotransmission-related pathology of MDD is state-related and mostly resolved during remission of MDD, and that remission is further associated with restructuring of intracellular components, structural elements and extracellular cell to cell contacts.

### CRH-, SST- and VIP-positive GABAergic cellular changes across MDD episodes and remission

The deconvolution of cell-type origin of gene expression obtained from gray matter samples (using independent sgACC single cell expression data) showed that a subpopulation of GABAergic inhibitory interneurons that express the CRH, SST and/or VIP neuropeptides are dynamically affected in synchrony with the episode and remission phases of MDD. Reduced CRH expression has been previously reported in subcortical brain regions, consistent with dysregulation of the hypothalamus pituitary adrenal stress axis in MDD, and we have previously reported reduced CRH expression in cortical layers as well (ref). Reduced SST expression has been reported by our group and others in several independent cohorts (refs). Reports on altered VIP expression in MDD are sparser (ref). Importantly, VIP, CRH and SST are markers of GABAergic interneuron subtypes that display overlapping patterns of expression, and that directly or indirectly inhibit pyramidal cell dendrites. These results are therefore consistent with a state-dependent MDD-related pathology that affects the regulation of excitatory input onto pyramidal cells, suggesting altered information processing in MDD (ref). At the sgACC regional level, reduced dendritic inhibition may alter the excitation inhibition balance and contribute to the elevated activity of this brain region during MDD episodes (refs).

The cellular deconvolution approach suggests variable proportions of a specific cell type between MDD episodes and remissions (Fig.3 D-E), but one could question how this can occur for interneurons, post-mitotic cells that seldom divide in the adult brain. Deconvolution analysis assumes that the gene expression of each cell type is linearly additive, making its contribution proportional to its fraction in bulk. Gene expression however, largely dependents on other cell-level covariate, particularly cell size and function. For instance, we previously showed a reduced SST expression per cell in the sgACC of MDD subjects, rather than reduced SST-positive cell numbers (Refs). The association of change in cell size or of expression per cell for CRH, SST and VIP positive interneurons is also supported by facts that normalizing for putative cell-type differences overall enhanced the MDD-related significance of cell-to-cell signaling and neurotransmission, and that multiple genes implicated in cell structure (hence volume and function) are affected here in phase with MDD episode and remission states.

### A putative causal role for immune function and inflammation in upper cortical layer cells in MDD

Bayesian network analysis is an approach that uses changes in association probabilities between gene modules across multiple network iterations to deduct potential causal flow. Applied to the dataset across control and MDD cohorts, this analysis suggests that activation of immune function, inflammation and oxidative stress originating from or affecting inhibitory neurons, glial, endothelial and immune cells in upper cortical layers may have causal role in MDD across episode and remission phases. The collective results may point to sustained immune activation, combined with other cellular stressors (oxidative stress, inflammation), whether of intrinsic or external origin, and implicating glial and endothelial cells, in turn affecting inhibitory neurons located in upper cortical layers and regulating processing of cortical information. This collective pathology is maintained throughout the disease trajectory but appears to wax and wane with episodes and remission, as suggested by phasic changes in CRH-/VIP-/SST-neurons, reversal of neuronal structure during remission, shift from innate and adaptive immune activation during episodes receding to innate immunity only during remission, and paralleled by bioenergetics and cellular structure phase-dependent changes. In this way, the model suggests that the trait pathology associated with immune/inflammation increases the biological vulnerability of the sgACC, setting it up for relapse into episode state-like pathology, together demonstrating a novel type of plasticity associated with MDD.

It is intriguing that the gene expression profile of the Bayesian network gene module most associated with MDD can be antagonized or mimicked in cell lines by drugs and ligands that target the dopamine or monoamine systems, consistent with the clinic roles of these neurotransmitter systems in diseases and therapeutics. It is surprising that this information can be obtained from acute drug exposure in simple cell-based systems that lack the complexity of the brain and the timeframe of disease trajectory. This may reflect the fact that the complex interplays of multiple biological events across time and cell systems in MDD may recruit a unique combination of cellular processes, as identified by correlated gene coexpression patterns between complex modules obtained in combined gray matter tissue across disease states and those induced by specific compounds on simplified cell systems. This is also consistent with novel therapeutic or pro-disease targets suggested by this approach, which belong to complex biological regulators, such as epigenetic, nuclear receptors and protein modification regulators, rather than single target molecules. Importantly, these findings provide an independent confirmation of the validity of the causal gene modules and biological pathways identified in this study. Moreover, the integrated approach applied in this study provides a framework and putative temporal sequence to assemble known hypotheses of MDD, namely suggesting immune- and cellular stress-related phenomenon as causal and upstream in series of event contributing to the state-dependent plasticity affecting excitatory and inhibitory neuron and transmission, as well as serotonin, glutamate and GABA dysfunction.

### Limitations

These studies were performed in cross-sectional cohorts, and results suggesting causal links and sequence of events should be interpreted as hypothesis generating, to be tested in future studies. These studies were aimed at identifying broad biological events rather than specificities associated with demographic (sex, age) or clinical (antidepressant use, suicide). Finally, the unique feature of the current cohorts (remission and episodes) and availability of samples precluded the validation of the state-dependent aspects of the current results in independent cohort. Instead we have used available genomic datasets to provide complementary or additional supporting evidence for aspects of the current results.

## MATERIAL AND METHODS

### Human postmortem brain samples

Postmortem brain samples were collected during routine autopsies performed at the Allegheny County Medical Examiner’s Office with procedures approved by the University of Pittsburgh committee for oversight of study and clinical training involving the dead. Consensus DSM-IV diagnoses were made by an independent committee of experienced clinicians using information from structured interviews with family members, clinical records, toxicology results, and standardized psychological autopsies. A similar approach was employed to confirm the absence of a psychiatric diagnosis in comparison subjects. The MDD cohorts were carefully matched with controls to ensure that they did not differ in mean age, postmortem interval (PMI), brain pH or RNA integrity number (RIN). Ninety samples including 20 control subjects, 20 subjects in first MDD episode, 15 in remission after a single episode, 20 in recurrent episode and 15 in remission after recurring episodes (fig.1, table S1). Samples comprising all six cortical layers were collected from coronal sections, as previously described *(42)*. Samples from the same cohort had been previously used in a large-scale proteomic study *(20)*.

### RNA extraction and sequencing library preparation

Total RNA was extracted from the sample homogenates using RNeasy Mini kit (Qiagen, Cat.No.74104) with in-column DNAse treatment using RNAase-Free DNase (Qiagen, Cat.No.79254). Sequencing libraries were prepared using SMARTer Stranded Total RNA-seq kit (Clontech Laboratories, Cat. No. 634876). All steps involved were performed according to manufacturer’s protocol.

### Sequencing and data generation

Pooled libraries were sequenced in illumina HiSeq2500 to generate 2 × 100 paired end reads, which were aligned to human reference genome GRCh38 provided by Ensembl using HISAT2 aligner. Count data were then generated for the reads aligned to exons and transcripts using the GenomicFeature and GenomicAlignments packages in R and gene model (GTF file) provided by Ensembl. 9.7 million reads were obtained per sample, aligning to 21,684 genes.

### Experimental contrasts and differential expression analysis

After removing low expressed genes (mean row sum <= 5 counts across all 90 samples), 21,684 genes were included in differential expression analysis. The experimental design allowed the following contrasts to be examined using the DESeq2 R package: 1) **MDD-all**: all MDD cohorts versus Controls, 2) **MDD-episode:** two MDD episode cohorts versus Controls (E_n_|C), 3) **MDD-remission:** two remission cohorts versus controls, 4) **Episode/Remission:** two MDD episode cohorts versus two remission cohorts (E_n_|R_n_), 5) **Progressive Episode**: monotonic increase or decrease across control and episode groups; C→E_1_→R_R_, 6) **Progressive Remission**: monotonic increase or decrease across controls and remission groups; C→R_1_→R_R_) and 7) **MDD-phasic**: capturing gene expression patterns coupled to the phasic changes across groups. The curve fitting for MDD-phasic was performed in two steps in R. For the first step. pData$MDD-Phasic = sin (-pi/2 + 2 * pi/2 * pData$group) adds the column named MDD-phasic to the sample phenotype data used to model the sinusoidal nature of the cohort. Here -pi/2 and pi/2 captures the down and up phase respectively. 2 denotes the step ensuring the model searches for alternate down and up pattern. “pData” denotes phenotype data table and group denotes the cohort where the order is preserved as C→ E_1_→R_1_→E_R_→R_R._ The second step involved using the MDD-phasic column in the phenotype data as contrast (similar to other contrast) for finding genes associated with it. For all contrasts, Likelihood ratio test was employed with a full design of ∼Age+Sex+PMI+Contrast and a reduced design of ∼Age+Sex+PMI to remove the effect of age, sex and postmortem interval (PMI). A p-value threshold of 0.05 was used.

### Pathway enrichment analysis

Biological pathways affected in different contrasts were determined using gene set enrichment analysis (GSEA) *(43)*. The 21,684 genes ranked by Wald statistic were tested against the three Gene Ontologies (GO): Biological Process (GOBP), Molecular Function (GOMF) and Cellular Component (GOCC). Updated gene set (pathway) lists were obtained from the Bader lab (http://download.baderlab.org/EM_Genesets/). To compare the effect of MDD pathology across different contrasts, the normalized enrichment score of significant pathways (P-value<0.05, q-value<0.25) was used to generate heatmaps.

A focused analysis of *a priori* functional themes was performed to better identify the character of the biological changes in the enrichment results. Specifically, we selected significantly enriched pathways which either 1) contained the name of the functional theme based on a text-based query or 2) were identified as child terms (nested pathways) of pre-selected parent terms which represent the theme.

### Hypergeometric analysis

To look for significant overlap between gene sets we performed hypergeometric overlap using Geneoverlap package in R. Background of 21196 genes and significance cutoff of p-value < 0.05 was used for all the analysis.

### Quantitative polymerase chain reaction (qPCR)

Differentially expressed (DE) genes belonging to a leading-edge subset (core set of transcripts that accounts for the enrichment signal) in our enrichment analysis were used to validate the differential expression results using qPCR. Top 5 samples representing either control or diseased state were selected based on their expression profile. Total RNA (same as used for generating the sequencing libraries) were reverse transcribed to cDNA using PrimeScript RT Master Mix (TaKaRa). cDNA, primers, and TB Green Premix ExTaq (Tli RNaseH Plus) (TaKaRa) were mixed in 96-well PCR plate, and qPCR was performed in triplicate using CFX96 Real-Time System (Bio-Rad). Results were normalized to GAPDH internal control. Primers used are described in Supplementary Table 2.

### Cellular Deconvolution analysis

To estimate the cell type proportion and identify putative disease-associated cellular differences, we adapted the deconvolution analysis described by Baron et. al. *(44)* and implemented in bscqsc package in R. The analysis consists of five steps. **A)** <u>Identifying cell-type-specific marker-genes</u> Using the single nucleus data set for Anterior cingulate cortex available form Allen brain atlas, we first identified cluster of cell types (fig.3A) and markers specific to each cluster (table. S9) using SEURAT package in R for single cell analysis *(45)*. In order to segregate cell clusters based on subtle differences in expression, the resolution parameter was set to 1.2 **B)** Building the reference basis matrix of marker-genes (fig.3D): This matrix contains the highly discriminatory marker-gene expression of each cell-type cluster averaged across all the cells of a given cell type cluster. For a given cluster, we considered a gene specific to a cluster only when it showed >= 3-fold difference in expression when compared to its expression in other cell clusters. **C)** Estimating proportions: The resulting reference matrix (fig.3D) was used to estimate cell proportion in different cohorts and contrast (fig.3E) using Support Vector Regression implemented by CIBERSORT package in R *(46)*. **D)** <u>Adjusting the bulk tissue gene expression differences for proportion</u>: We statistically regress out the effect of a cell types cluster (cluster 8) which showed the significant change in proportion in Episode/Remission contrast. This was implemented by expanding the design-model used for finding differential expression between episode and remission cohort by incorporating the estimated cell-type proportion (from step C) in the design matrix. Note that this step allows for differential expression analysis independent of variation introduced by cell proportion. Comparing the differentially expressed gene with and without the adjusted cell-type proportion can be used to identify genes which show increased statistical significance (i.e. further reduced p-value) after removing the effect of cells showing highest change in proportion for a given contrast. These set of genes and the pathways associated with them can be considered as those influenced (thus consequential). **E)** <u>Finding cell-type specific differential expression</u>: As described previously *(47)* and implemented in the csSAM (a package in R) we use the two separate differential expression analyses, (i.e. with and without regressing the cell proportion) to fit a linear model that estimates interaction term between cell proportion and contrast to find cell type cluster specific differential expression

### Bayesian Network analysis

To find probabilistic causal associations of disease states with gene co-expression modules representing different biological themes we used weighted gene coexpression network analysis (WGCNA) and Pigengene package in R. First, we used consensus WGCNA to generate co-expression modules common across samples and compared them between control and disease states. For each cohort, count data normalized based on size factor (using DESeq2 package in R) was used to create a matrix of pairwise Pearson correlations between genes, which was then transformed to a signed adjacency matrix using power β=12. To calculate the interconnectedness between genes, which defines “modules”, we derived the topological overlap, which gives biologically meaningful measurement of similarity between two genes, based on their co-expression relationship with all other genes. We then identified consensus modules between the control and the diseased cohorts using blockwiseConsensusModules function with consensusQuantile setting set to 0.50. We obtained 22-consensus modules with numeric label 1 to 22 and color labels for all. Label 0 and color grey was assigned to genes not assigned to any module and was removed from further analysis. The identified modules were functionally characterized using all three Gene Ontologies and top pathways with Bonferroni corrected p-value < 0.05 were used to label the modules.

The first principal component (the eigengene) summarizes a given module by accounting for the maximum variability of all its constituent genes. It is used as a proxy for the overall gene expression of this module and to identify mechanisms associated with disease states.

Using the Pigengene R package, module eigengenes were used as random variables to train a Bayesian network that modeled the probabilistic dependencies between all modules. Additionally, experimental group was included as a categorical variable (disease) to simplify the inference. The disease node works as a source and has no parent terms above it. This is implemented by blacklisting all the incoming edges to the disease node. To search for an optimal Bayesian network fitting our data, we performed 1000 permutations (i.e. 1000 networks) with default parameters in the Pigengen package. Edges that appeared in greater than equal to 30% of the networks were used to construct the network.

### Connectivity map (Cmap) analysis

Cmap contains expression profiles induced by ∼19,000 different small molecules (perturbagens) in ∼77 different cell lines *(48)*.Genes in module 10 were submitted as query in the Cmap API (https://clue.io/query). The output file (cs_nlx476251.gct) containing the raw weighted connectivity scores for all molecules, or perturbagens, tested in different experimental conditions and cell types was used for further analysis. A connectivity scores summarizes the similarity between the query and signature profile of a perturbagen based on Kolmogorov-Smirnov enrichment statistics. To compare an observed connectivity score to all others in the database we calculate the percentile score, referred as “tau”. The weighted connectivity score ranges from -1 to +1 and accordingly the tau ranged from -100 to +100. Negative and positive scores indicate dissimilarity and similarity between query and perturbagen signatures, respectively. In this study we restricted our search space to all perturbagen tested in 9 hallmark cancer cell lines (A375, A549, HA1E, HCC515, HEPG2, HT29, MCF7, PC3, VACP) frequently used in drug repositioning studies and three neuronal cell lines (NEU, NEC.KCL, NPC) available in the Cmap database. We summarized the multiple experimental conditions under which a given perturbagen was tested in a given cell line separately for each cancer and neuronal cell lines based on the largest effect size magnitude between 33^rd^ and 67^th^ percentile. To select the top perturbagen candidate, we used the rank ordered row sum of tau score, separately for cancer and neuronal cell lines.

### Drug target identification

Using the available two-dimensional (2D) structures (http://lincsportal.ccs.miami.edu/SmallMolecules/catalog) for top drugs associated with Module M10 and M16 molecules we searched for protein target and target class (protein family to which the target belongs) using SwissTargetPrediction, a web tool based working on structure function relationship of drugs and biomolecule. The frequency of different target class associated with the top 20 therapeutic drugs predicted based on neuronal cell lines was used to estimate the difference between drugs associated with M10 and M16 using chi-square test of independence. A drug can potentially dock to many protein targets of different target class and can hinder the calculation target class frequency. To avoid this issue we estimated the frequency based on the first predicted class target class for all the drugs.

## Notes

### Competing Interest Statement

The authors have declared no competing interest.

